# Rab-mediated trafficking in the secondary cells of Drosophila male accessory glands and its role in fecundity

**DOI:** 10.1101/266353

**Authors:** E. Prince, M. Brankatschk, B. Kroeger, D. Gligorov, C. Wilson, S. Eaton, F. Karch, R.K. Maeda

## Abstract

It is known that the male seminal fluid contains factors that affect female post-mating behavior and physiology. In *Drosophila,* most of these factors are secreted by the two epithelial cell types that make up the male accessory gland: the main and secondary cells. Although secondary cells represent only 4% of the cells of the accessory gland, their contribution to the male seminal fluid is essential for sustaining the female post-mating response. To better understand the function of the secondary cells, here we investigate their molecular organization, particularly with respect to the intracellular membrane transport machinery. We determined that large vacuole-like structures found in the secondary cells are trafficking hubs labeled by Rab6, 7, 11 and 19. Furthermore, these cell-specific organelles are essential for the long-term post-mating behavior of females and that their formation is directly dependent upon Rab6. Our discovery adds to our understanding of Rab proteins function in secretory cells. We have created an online, open-access imaging resource as a valuable tool for the intracellular membrane and protein traffic community.

## Summary statements

Secondary cells employ a previously unreported [Rab6-dependent] secretory system

## Introduction

Due to limited resources, sexual reproduction often leads to males having to compete to produce offspring in the succeeding generation (Agrawal, 2001; Andersson, 1994; Birkhead and Møller, 1998; Darwin, 1871; Smith, 2012). Thus, many organisms have developed methods to ensure the propagation of an individual’s genome at the expense of rivals (Clutton-Brock, 1989). For example, male polar bears often kill the offspring of rival males in order to favor the propagation of their own offspring (Lukas and Huchard, 2014). In *Drosophila melanogaster*, a more indirect “mate-guarding” strategy is used. The seminal fluid (SF) of *Drosophila* males contains factors, called seminal fluid proteins (SFPs), that are deposited into the female through mating (Wolfner, 1997, 2002). These factors influence the physiology and behavior of mated females and, in that way, favor the reproductive success of a mating male (Avila et al., 2011; Wolfner, 1997, 2002). The male-induced changes in mated females are called the <u>P</u>ost-<u>M</u>ating <u>R</u>esponse (PMR). Some characteristics of the PMR are: 1. decreased mating receptivity (Grillet et al., 2006; Gromko et al., 1984),2. reduced life span (Wigby and Chapman, 2005), 3. sperm storage (Adams and Wolfner, 2007; Ravi Ram and Wolfner, 2007; Wong et al., 2008), 4. increased ovulation (Heifetz et al., 2005; Rubinstein and Wolfner, 2013), 5. changed feeding behavior (Hussain et al., 2016) and 6. gut remodeling (Reiff et al., 2015). Mate-guarding strategies are also described for mammals but the mechanistic principles are less well understood. For instance, changes are reported for the ovulation frequency and immune response activity of mated females (Bromfield, 2016; Bromfield et al., 2014).

While in mammals, SFPs are mostly produced in the prostate gland, the seminal vesicles and the bulbourethral gland, in *Drosophila* these proteins are primarily produced by a single gland called the male accessory gland (AG). The *Drosophila* AG is a two-lobed structure, made of two types of bi-nucleated and secretory cell types arranged in a cellular monolayer surrounding a central lumen and surrounded by a layer of muscle cells. The two types of secretory cells have been named the main cells (MCs) and the secondary cells (SCs). The polygonally-shaped MCs make up 96% of the secretory cells of the gland and are known to produce the vast majority of the SFPs (Chapman and Davies, 2004; Kalb et al., 1993). The remaining 4% of secretory cells are the SCs, which are located only at the distal tip of the glands, interspersed between MCs; they are much larger, spherically-shaped and contain a number of large vacuole-like compartments (VLCs) (Bairati, 1968; Bertram et al., 1992; Gligorov et al., 2013). These cells, like the MCs, are in direct contact with the glandular lumen and are able to contribute to the seminal fluid (Bairati, 1968; Corrigan et al., 2014; Gligorov et al., 2013; Leiblich et al., 2012; Minami et al., 2012; Redhai et al., 2016; Sitnik et al., 2016). Recent findings show that the SCs are not crucial for initiating early PMR behaviors. Instead SC products play a critical role in extending the female PMR past the initial 24-48 hours after mating (Corrigan et al., 2014; Gligorov et al., 2013; Minami et al., 2012; Redhai et al., 2016; Sitnik et al., 2016). Interestingly, mutations that affect SC differentiation also impede the formation of their VLCs. VLCs are prominent membrane-bound organelles containing a large internal space. In mammals, VLCs have been implicated in different intracellular trafficking pathways such as endocytosis (Wada, 2013) or secretion (Jamieson and Palade, 1971).

Intracellular membrane and protein traffic is regulated by a family of membrane-associated, small GTPases called Rabs (Ras like bovine proteins). Since Rabs control individual trafficking sub-steps, these proteins are suitable to identify cellular membrane compartments (Bhuin and Roy, 2014; Hutagalung and Novick, 2011). Recently, a collection of YFP-tagged *rab* alleles was established in *Drosophila* (Dunst et al., 2015). This Rab library allows *in vivo* tracking and manipulation of the Rab proteins in any given cell type (Dunst et al., 2015).

Here, we use the Rab library to screen for the expression and localization of all *Drosophila* Rab proteins in the *Drosophila* AG. Focusing on the SCs, we show that Rab6, 7, 11 and 19 define four different VLC populations. This extends previous studies that showed that there were at least two different subclasses of VLCs using numerous intracellular markers (Bairati, 1968; Leiblich et al., 2012; Corrigan et al., 2014). Furthermore, we track the development of VLCs over the first few days after male eclosion and find that all founding VLCs that we detect are Rab6-positive, while Rab7-, Rab11- and Rab19-positive compartments appear later in adulthood, suggesting they may be Rab6-dependent. Consistent with this finding, the genetic reduction of Rab6 prevents the formation of all VLC classes. However, the loss of Rab7 and 11 (Corrigan et al., 2014; Redhai et al., 2016), but not Rab19, also results in the loss of specific VLCs and change female PMR behaviors. Finally, we have established an online-based imaging platform (https://flyrabag.genev.unige.ch). This resource provides annotations based on a defined vocabulary for each expressed Rab protein in AG and allows 3D localization tracking down to subcellular resolutions. For the first time, the membrane/protein transport machinery of the AGs is comprehensively charted and our findings add valuable knowledge to the existing model describing the SC secretion system (Bairati, 1968; Bertram et al., 1992; Corrigan et al., 2014; Kalb et al., 1993; Leiblich et al., 2012; Redhai et al., 2016; Sitnik et al., 2016; Wilson et al., 2017)

## Results

### The basic morphology of the accessory gland epithelium

An ultrastructural study of the gland performed by Bairati in 1968 (Bairati, 1968) suggested that both AG cell types are polarized and secretory in nature. To confirm this notion, we examined the cellular localization of the canonical cell polarity markers, DE-Cadherin (DCAD, marking apical adherence junctions) and Disc-Large (Dlg, marking basolateral membrane), as well as the F-actin staining molecules, Phalloidin and Lifeactin (Schnorrer, 2011), in *Drosophila* AGs. Confirming the findings of Bairati, we find that the cells of the AG are indeed polarized, with their apical surface facing the central lumen (Figs 1B-1D). One striking characteristic of the cellular monolayer making up the AG regards the SCs. The SCs display a distinct round shape and seem to extrude out from the uniform sheet of MCs into the luminal space (Figs 1A-1B). However, even with this extrusion, they do not display a large exposed surface to the luminal fluid, as the MCs surrounding the SCs seem to extend over much of their surface to restrict contact with the lumen (Figs 1A & 1C) (Leiblich et al., 2012). This spreading of the MCs over the SCs also results in a large contact zone between the two cell types. Furthermore, we find a dense F-actin network concentrated along the apical surface of the SCs, with actin-filled membrane protrusions extending into the luminal space (Fig1D) (Bairati, 1968). This apical enrichment of F-actin is reminiscent of other secretory gland cells (eg. salivary gland cells) and may reflect the secretory nature of SCs (Dunst et al., 2015; Myat and Andrew, 2002).

**Figure 1.**
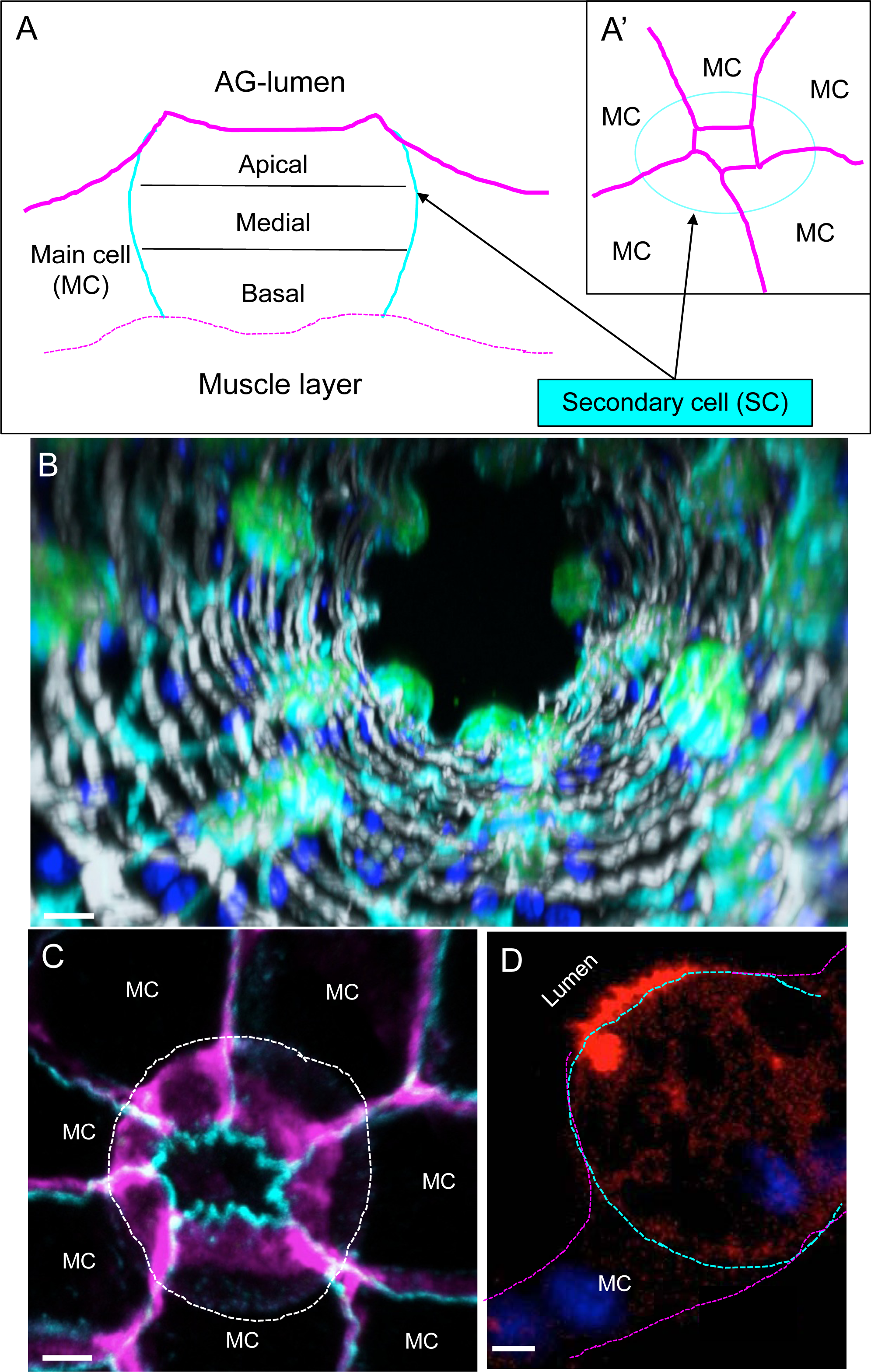
Organization of the Secondary cells. (A and A’) Schematic depiction of a secondary (SC, light blue) and flanking main cells (MCs, magenta). SC cells are embedded in MCs. We divided SCs into three zones: Apical, Medial and Basal (A, sagittal view). The apical, luminal contact zone of SCs is very small (A’, top-view, magenta). (B) Confocal images from the distal tip of an AG are assembled into a 3D projection down the long axis of the gland. GFP (green) expressing SCs are probed for Dlg (cyan), F-actin (light-grey) and DAPI (dark blue). Scale bar = 15μm. (C) Compressed confocal stack (15µm from apical to basal) shows a SC (outlined by a white dashed line) probed for DCAD (cyan) and Dlg (magenta). Surrounding main cells (MCs) are indicated, scale bar = 5μm. (D) A confocal slice along the apical-basal axis of a representative SC (outlined with a dashed line) expressing the F-actin marker, Lifeactin-Ruby (red), and stained with DAPI (blue). The apical side of the cell is at the top left and the basal side is at the bottom right. Note the enrichment of actin filaments at the apical membrane that is facing the AG lumen. Scale bar = 5μm.

### Secondary cells have different vacuole-like compartments

Although VLCs are prominent in SCs (Bairati, 1968; Bertram et al., 1992; Gligorov et al., 2013), their molecular organization and function remains elusive. We hypothesized that VLCs could be a single trafficking compartment required for the efficient secretion of SFPs. To study the role of VLCs in membrane trafficking, we screened all expressed Rab proteins using the YRab-library (Dunst et al., 2015) (http://rablibrary.mpi-cbg.de/). To annotate the localization of each expressed Rab protein we used a defined terminology (see Materials & Methods) and show multiple representative original confocal data sets at our CATMAID-based website (https://flyrabag.genev.unige.ch). In this way, users are able to navigate and track Rab compartments at subcellular resolution. Here, we will primarily focus on the Rab localization patterns in the SCs. Overall, we find that 16 of Rabs are expressed in SCs and that 4 of these Rabs are associated with VLCs: Rab6, 7, 11 and 19 (Fig 2, https://flyrabag.genev.unige.ch).

**Figure 2.**
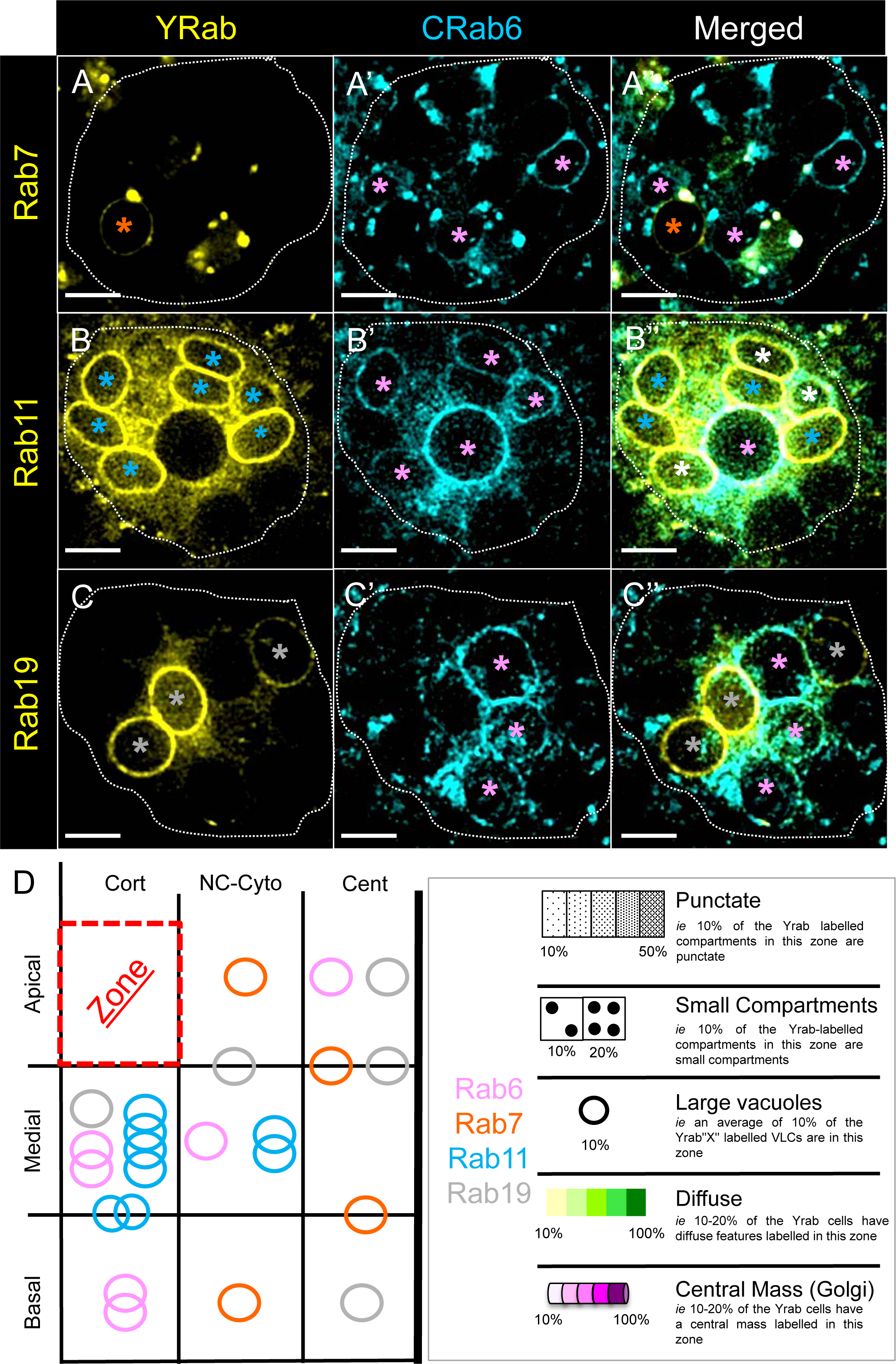
The Rabs associated with VLCs. (A-C’’) Shown are individual wide-field florescence microscopy slices (0.8µm slices) of SCs at three levels along the apical-basal axis (A-A’’ Apical; B-B” Basal; C-C” Apical) from *Crab6; Yrab7*, *Crab6; Yrab11* or *Crab6; Yrab19* flies. Non-fixed AGs were visualized for YFP (yellow; A, B, C) and CFP (cyan; A’, B’, C’) fluorescence. Asterisks indicate labeled VLCs (orange, Yrab7; pink, Crab6; light blue, Yrab11; grey, Yrab19; white, for VLCs in the merged images that co express two Rabs), scale bars = 5µm. (D) Plotted is the relative position and abundance of VLCs labelled by one Rab in SC cells; color code and symbols are annotated in the legend (right box). Each circle indicates approximate proportion of the vacuoles (based on the average proportion of vacuoles labeled per cell) labeled by the particular rab in the indicated zone (Rab6, n_cell_=19, n_VLCs_/cell=6.37± 2.69; Rab7, n_cell_=8, n_VLCs_/cell=4.38 ± 3.34; Rab11, n_cell_=5, n_VLCs_/cell=9.4 ± 2.7; Rab19, n_cell_=13, n_VLCs_/cell=6.08 ± 2.93). Abbreviations: Cort = Cortical; NC-Cyto = Non-Central Cytoplasmic; Cent = Central. A key to the symbols used in all schematic representations of Rab localization is present next to this plot (not all symbols are used in this plot). The percentage indications for each structure are defined as follows: for the punctate and small compartments the percentages indicate that the particular structure in that zone makes up, on average, the indicated percentage of the total Rab labeling in a cell (average of percentages from n cells). For the diffuse and central mass, the percentages indicate the percentage of cells (n) with that structure in that zone (Rab6, n_cell_=19; Rab7, n_cell_=8; Rab11, n_cell_=5; Rab19, n_cell_=13).

### Rab6 is associated with the *trans-*Golgi Network and VLCs

Rab6 is known to localize within the *trans*-Golgi network (TGN) and to regulate protein and membrane traffic from the Golgi organelle to other membrane targets (Dunst et al., 2015; Iwanami et al., 2016; Satoh et al., 2016; Schotman and Rabouille, 2009). To test if VLCs are interconnected to the TGN, we probed *Yrab6* glands together with a battery of known Golgi-markers (Dunst et al., 2015; Harris, 2016; Rikhy and Lippincott-Schwartz, 2010). As expected, Rab6 is associated with the Golgi organelle in SCs (Fig 3D). However, the appearance of the Golgi in SCs (Fig. S1A) is very different from the Golgi organelle in other cells (eg. MCs) (Bairati, 1968). In most *Drosophila* cell types, multiple Golgi units are dispersed and their build-up is primitive, consisting of a single *cis-*Golgi and *trans-*Golgi membrane sheet (Kondylis and Rabouille, 2003, 2009; Rabouille et al., 1999; Ripoche et al., 1994; Schotman and Rabouille, 2009). In SCs, we find that the Golgi forms a central, extended structure within the basal-medial area of the cell (Fig. S1A).

**Figure 3.**
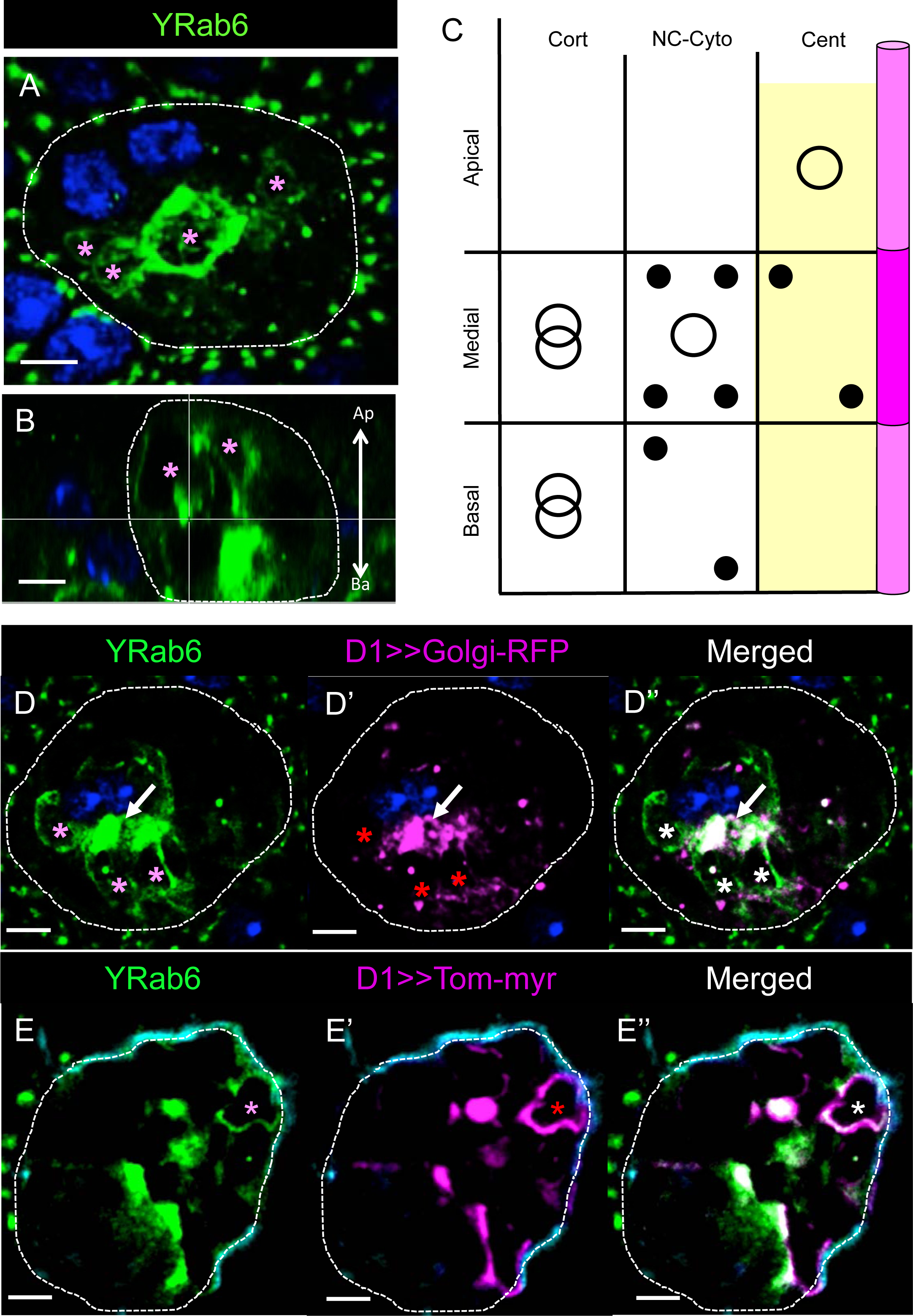
Organization of Rab6 membranes in SCs. (A, B) Shown is a Z-projection (A, 15µm; Medial; top-view) and a confocal reconstruction of a sagittal section from the same confocal stack (B, 0.8µm; sagittal-view) of a Yrab6 SC probed for DAPI (blue) and YFP (green). Pink asterisks indicate Yrab6-VLCs, scale bars = 5µm, arrow shows the apical-basal cell orientation (B). (C) Plotted is the position and relative abundance of Rab6-positive compartments. Color code and symbols are explained in Figure 2D (right box and legend) (n_cell_=19). (D-E”) Shown are confocal slices of single SCs (0.8µm thick slices; D-D’’ Medial; E-E” Medial) from *Yrab6; D1>>golgiRFP* (D-D”) and *Yrab6; D1>>tomato^myr^* (E-E”) flies. AGs were probed for YFP (green), DAPI (dark blue) and RFP/Tomato (magenta). Asterisks indicate labelled VLCs (pink, Yrab6; red, Tomato/RFP markers; white and white arrows, for colocalization), scale bars = 5µm.

The VLCs bound by Rab6 (n_cell_= 19; n_VLCs_/cell=6.37± 2.69), however, display no Golgi signature (Fig 3D). Rab6-VLCs appear mainly in two areas: most are found in the basal-to-medial part of the cell along the plasma membrane, while other VLCs appear apically localized in the non-central (NC)-cytoplasmic and central regions of the cell (Figs 3A-3C, and Materials and Methods for location terminology). The distribution of Rab6 in SCs is somewhat similar to salivary gland (SG) cells, where Rab6 is found on Golgi but also on non-Golgi compartments (Dunst et al., 2015). Nevertheless, the extreme enlargement of all Rab6 compartments in the SCs and the more-differentiated morphology of the Golgi, point to extensive membrane/protein transport processes (Liu and Storrie, 2012, 2015; Rodriguez-Boulan and Macara, 2014; Rodriguez-Boulan et al., 2005).

To test if Rab6-VLCs are involved in protein secretion, we expressed myristoylated fluorescent protein, Tomato (Tomato^myr^) (Pfeiffer, 2010), in *Yrab6* SCs. These results show that the cortical and NC-cytoplasmic Rab6-positive compartments are used to transport this reporter protein (Fig 3E). Thus, Rab6-VLCs are a route for secreted proteins.

### Rab19-labeled VLCs are dependent on *Rab7*

In SGs, Rab19 is found exclusively associated with the apical membrane (Dunst et al., 2015). Although the biological role of Rab19 is poorly understood, Rab19 has been suggested to be involved in apical secretion (Dunst et al., 2015). We found that in AGs, Rab19 is strongly expressed in SCs and associated with VLCs (Fig 4, n_cell_=13, n_VLCs_/cell=6.08 ± 2.93). Surprisingly, none of the Rab19-VLCs shows Rab6 co-localization (Fig 2C) and only a small proportion of Tomato^myr^ is trafficked in these compartments (Fig 4E). Of note, a small fraction of Rab19 is localized on Golgi membrane domains (Fig 4D). Based on these results, Rab19 may regulate another VLC traffic route, distinct from Rab6 transport.

**Figure 4.**
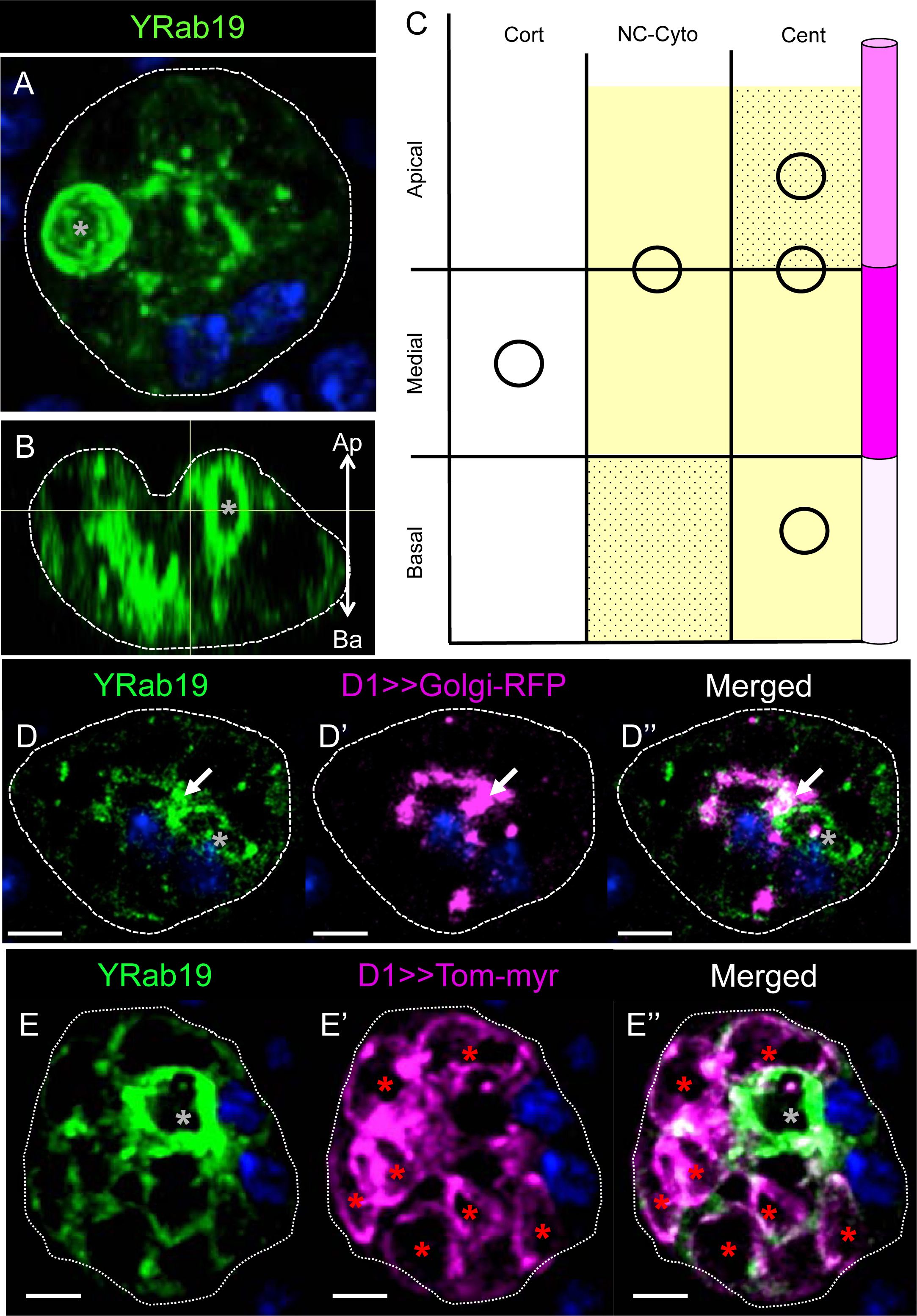
Organization of Rab19 membranes in SCs. (A, B) Shown is a Z-projection (A, 15µm; Medial; top-view) and a confocal reconstruction of a sagittal section from the same confocal stack (B, 0.8µm; sagittal-view) of a Yrab19 SC probed for DAPI (blue) and YFP (green). Grey asterisks indicate Yrab19-VLCs, scale bars = 5µm, arrow shows the apical-basal cell orientation (B). (C) Plotted is the position and relative abundance of Rab19-positive compartments. Color code and symbols are explained in Figure 2D (right box and legend) (n_cell_=13). (D-E”) Shown are confocal slices of single SCs (0.8µm thick sections; D-D’’ Apical; E-E” Apical) from *Yrab19; D1>>golgiRFP* (D-D”), and *Yrab19; D1>>tomato^myr^* (E-E”) flies. AGs were probed for YFP (green), DAPI (dark blue) and RFP/Tomato (magenta). Asterisks indicate labelled VLCs (grey, Yrab19; red, Tomato/RFP markers; white, for colocalization), scale bars = 5µm.

Interestingly, we also found Rab7 associated with Rab6-negative VLCs in SCs (n_cell_=8, n_VLCs_/cell=4.38 ± 3.34). Rab7 is known to regulate late endosomal traffic (Fig. S2) and is enriched on lysosomes ((Bucci et al., 2000; Corrigan et al., 2014; Meresse et al., 1995). In addition, Rab7 is found on ER exit sites where it co-localizes with Rab1 (Caviglia et al., 2017). The association of Rab7 with ER/*cis*-Golgi membranes may indicate the formation of specialized Rab7 compartments that enter non-canonical trafficking routes (Caviglia et al., 2016). Thus, we speculated that Rab7 membranes might show an interrelation with Rab19-VLCs. To test our idea, we knocked down *Rab7*. Astonishingly, loss of Rab7 results in the complete depletion of Rab19- (Fig 6D’’) but not Rab6-VLCs (Fig 6A’’). On the other hand, loss of Rab19 changes the appearance of Rab7-VLCs but is not required for their existence (Fig 6B’’’). We conclude, that Rab19 probably belongs to a Rab7-related endocytic traffic cascade in SCs.

### Rab11-VLCs partially overlap with Rab6 traffic

Rab11 is generally associated with recycling endosomes (Bhuin and Roy, 2011, 2014; Casanova et al., 1999; Hutagalung and Novick, 2011; Ullrich et al., 1996) and is known to contribute to SC secretory activity (Corrigan et al., 2014; Redhai et al., 2016). Rab11-VLCs are mainly located near the plasma membrane (cortical) around the basal/medial and medial part of the SCs (n_cell_=5, n_VLCs_/cell=9.4 ± 2.7). Rab11 also marks punctae, basally-to-medially enriched in the cytoplasmic compartment and punctae in the central area with an apical enrichment (Fig 5). Interestingly, Rab11 and Rab6 seem to be present on VLCs located in the same zone of the SCs (Figs 2B, 2D, 5D). To test if Rab6 and 11 label the same VLCs, we probed AGs that express both Yrab11 and Crab6 (like Yrab6 but marked with CFP). We find that Rab6 and Rab11 do coexist on some, but not all VLC membranes (Fig 2B), suggesting that both of these VLCs may be involved in the same secretion pathway. To test this, we expressed Tomato^myr^ in the SCs of *Crab6; Yrab11* males. Importantly, we find Tomato^myr^ protein in CRab6- and CRab6/YRab11-VLCs, suggesting that Rab11 is another traffic checkpoint in the Rab6-aligned secretion pathway (Fig 5E, (Iwanami et al., 2016)).

**Figure 5.**
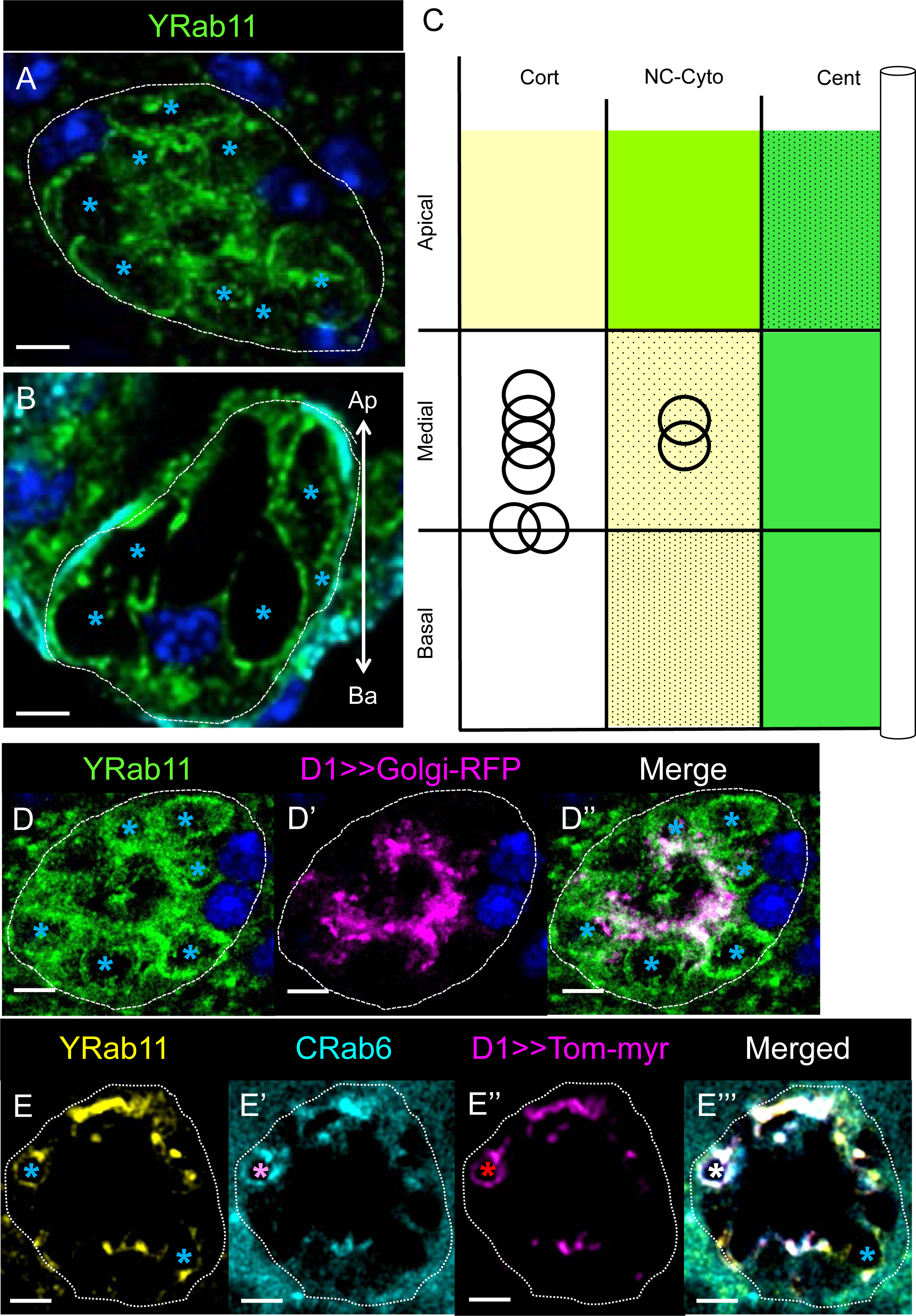
Organization of Rab11 membranes in SCs. (A, B) Shown is a Z-projection (A, 15µm; Medial; top-view) and a confocal reconstruction of a sagittal section from the same confocal stack (B, 0.8µm; sagittal-view) of a Yrab11 SC probed for DAPI (blue) and YFP (green). Light blue asterisks indicate Yrab11-VLCs, scale bars = 5µm, arrow shows the apical-basal cell orientation (B). (C) Plotted is the position and relative abundance of Rab11-positive compartments. Color code and symbols are explained in Figure 2D (right box and legend) (n_cell_=5). (D-E”) Shown are confocal slices of single SCs (0.8µm thick sections; D-D’’ Basal; E-E” Basal) from *Yrab11; D1>>golgiRFP* (D-D”) and *Crab6*; *Yrab11, D1>>tomato^myr^* (E-E’’’) flies. AGs were probed for YFP (green), CFP (cyan) and RFP/Tomato (magenta). Asterisks indicate labelled VLCs (light blue, Yrab11; red, Tomato marker; pink, Crab6; white, for colocalization), scale bars = 5µm. In (E’’’). A VLC marked by the Crab6, Yrab11 and *tomato^myr^* is indicated by a white asterisk (E’’’).

### VLC formation and the female long-term PMR are Rab6-dependent

Recent studies have shown that AGs require four days to reach full maturity/functionality (Ruhmann et al., 2016). To investigate if VLCs change their molecular identity in that developmental timeframe, we tracked the VLC formation in SCs over time (Fig. S3). Interestingly, one hour after eclosing, SCs show exclusively Rab6-VLCs. Later, at five hours post-eclosion, VLCs with Rab19 and VLCs with Rab11 identity become visible. It is only after three days that Rab7-VLCs appear. Over the next two days, the VLCs continue to grow in size and number until five days post-eclosion, when the cells seem to reach a stable structure. These results, taken together with the functional data (Ruhmann et al., 2016), correlate the development of the VLCs with AG functionality in newly eclosed adult males (Ruhmann et al., 2016).

The association of specific Rabs to maturing VLCs begs the question of whether or not the Rabs are directly required for the formation of the VLCs and SC functionality. Previously, we have shown that the maintenance of the PMR is impaired in mutants that lack VLCs filling their cytoplasm (Gligorov et al., 2013). Therefore, we chose to knock-down each candidate *Rab* individually in SCs and to assay VLC appearance and the female long-term PMR behavior. Strikingly, knocking down *Rab6* in the SCs leads to the disappearance of all VLCs in mature AGs (Figs 6A’-D’ and Fig. S4, S5). It is interesting to note that smaller vesicles marked by Rab7, Rab11 and Rab19 are still present (Figs 6B’-D’) and that the Golgi-RFP marker still marks elements of a central channel (Fig. S5). Furthermore, Tomato^myr^ production is not impaired in SCs lacking Rab6 and its membrane association indicates that Golgi post-translational modifications are still possible (data not shown). Thus, the loss of VLCs in the *Rab6* knockdown cannot simply be explained by either the absence of Rab7, 11 or 19 protein (Fig 6B’-D’, S4), or the complete lack of Golgi apparatus (Fig. S5) and may be the result of a Rab6-VLC maturation process.

**Figure 6.**
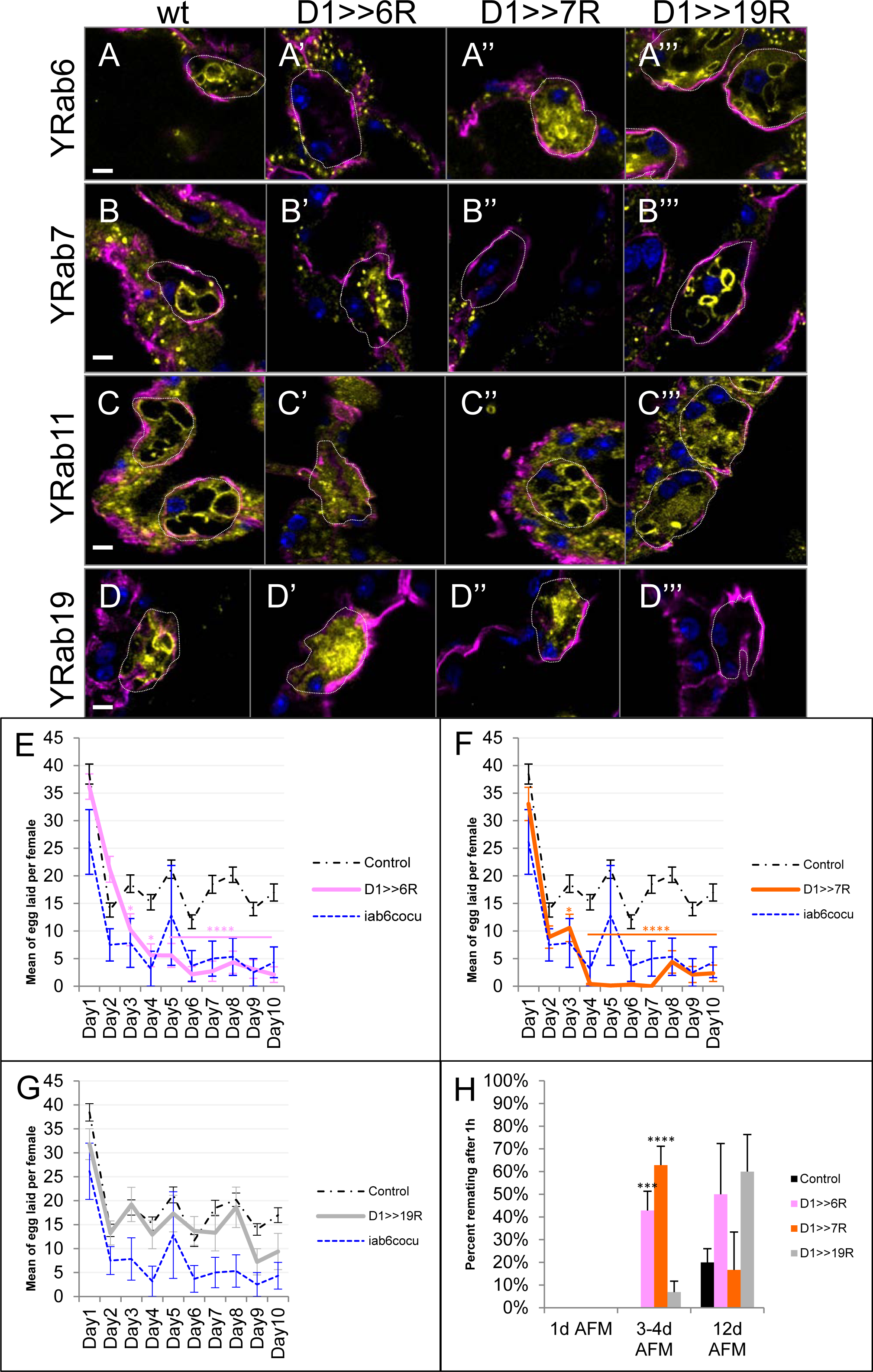
Rab6 is instructive for VLC formation and the female PMR. (A-D’’’) Shown are confocal slices (0.8µm thick sections; top-view; A-A’’’, Medial; B-B’’’ and DD’’’, Apical; C-C’’’, Basal) of SCs from *Yrab6* (A), *Yrab6;D1>>rab6^RNAi^* (A’), *Yrab6;D1>>rab7^RNAi^* (A”), *Yrab6;D1>>rab19^RNAi^* (A’’’), *Yrab7 (B), Yrab7;D1>>rab6^RNAi^* (B’), *Yrab7;D1>>rab7^RNAi^* (B”), *Yrab7;D1>>rab19^RNAi^* (B’’’), *Yrab11* (C), *Yrab11;D1>>rab6^RNAi^* (C’), *Yrab11;D1>>rab7^RNAi^* (C”), *Yrab11;D1>>rab19^RNAi^* (C’’’), *Yrab19* (D), *Yrab19;D1>>rab6^RNAi^* (D’), *Yrab19;D1>>rab7^RNAi^* (D”) and *Yrab19;D1>>rab19^RNAi^* (D’’’) *flies.* AGs were probed for YFP (yellow), Dlg (magenta) and DAPI (blue). Scale bars = 5µm. (E-G) Plotted are the number of eggs laid over a ten-day period for *wild-type* females mated with males of following genotypes: *D1>Gal4* and *Canton-S* (control, black broken line), *iab6^cocu^* (blue broken line), *D1>>rab6^RNAi^* (E, pink), *D1>>rab7^RNAi^* (F, orange), *D1>>rab19^RNAi^* (G, grey). Standard error of the mean is indicated. Statistics, *, p<0.05; **, p<0.01; ****p<0.00001; Mann-Whitney U test. (H) Shown is the re-mating frequency (in %) of *wild-type* females, previously mated with males of following genotypes: *D1>>GFP* (control, black), *D1>>rab6^RNAi^* (pink), *D1>>rab7^RNAi^* (orange), *D1>>rab19^RNAi^* (grey). The males used for the secondary matings are *wild-type*. AFM indicates time after the initial mating, error bars indicate standard error of the mean. Statistics, *, p<0.05; **, p<0.01; ****p<0.00001; Mann-Whitney U test.

The loss of Rab6 in SCs also results in a dramatic decrease in the long-term but not short-term PMR (Figs 6E, 6H). Although the PMR starts normally, the mating-induced egg-laying stimulation (Fig 6E) and the reduction of secondary mating receptivity (Fig 6H) is not sustained past the first two days post-mating. This is similar to the PMR seen in mates of *iab6^cocu^* mutant males who also lack SC VLCs ((Gligorov et al., 2013), Fig 6E). Although the systematic functional analysis of *Rab11* was prevented due to general lethality, we were able to test for the effects of *Rab7* and *Rab19* knockdown. Knockdown of either *Rab* did not affect the formation of Rab6- (Figs 6A’, 6D’) or Rab11-VLCs (Figs 6A’’’, 6D’’’). However, the absence of Rab7 in SCs prevents the formation of Rab19-VLCs and also changes the long-term PMR (Figs 6F, 6H). As the knockdown of *Rab19* does not affect the long-term PMR, we presume, that the effect of *Rab7* knockdown stems from a central endocytic block that impairs the general functionality of SCs (Corrigan et al., 2014). Following this interpretation, Rab19 may belong to an unrelated, more specialized trafficking pathway.

## Discussion

Recent findings regarding the principles of intracellular protein/membrane trafficking have shown that different cell types show a surprising versatility with regards to their usage of the intracellular transport machinery. Although the Rab protein family regulates intracellular traffic steps, across different cell classes (*i.e.* epithelia), the composition, organization and trafficking function of Rab proteins often seems incomparable (Caviglia et al., 2017; Dunst et al., 2015; Fu et al., 2017). Therefore, it is vital to chart and understand intracellular trafficking pathways in as many suitable model systems as possible to describe general transport principles, like continuous or pulsed secretion (Caviglia et al., 2017; Dunst et al., 2015; Fu et al., 2017; Iwanami et al., 2016; Redhai et al., 2016).

Here, we describe the protein trafficking pathway organization of the SCs of the *Drosophila* male AGs and lay down a molecular foundation for Rab-dependent transport routes in this cell type. We and others found that SCs are embedded in a monolayer of primary cells (MCs) and that their apical side faces the central gland lumen (Bairati, 1968; Corrigan et al., 2014; Leiblich et al., 2012; Redhai et al., 2016). However, it is interesting to note that the luminal membrane of SCs is highly restricted by the surrounding MCs and that there is a large apical SC/MC contact zone. These overlapping membranes may form an intercellular cavity and secreted proteins could be trapped between apical adherence and baso-lateral contact zones. Such a morphological feature is known to facilitate paracellular transport (Bökel et al., 2006; Marois et al., 2006; Wucherpfennig et al., 2003). If true, SCs are perfectly positioned to receive material from neighboring MCs and to secrete these products into the gland lumen. In support of this idea, it was reported that the SFP Ovulin is produced in MCs (Gligorov et al., 2013; Kalb et al., 1993) but is found in the SC VLCs. This finding implies that MC-produced Ovulin can be endocytosed by SCs for protein modification (Gligorov et al., 2013).

To investigate this possibility and others, we decided to describe the transport machinery of SCs with our main focus on the Rab proteins and the Golgi network. We found that most Rabs are expressed in SCs and we used a defined terminology to annotate their intracellular localization. Our data are presented in our open access online platform (https://flyrabag.genev.unige.ch) and the approach is complementary to an already published online resource for other *Drosophila* cell types (Dunst et al., 2015) (http://rablibrary.mpi-cbg.de/).

One predominant intracellular compartment in SCs are VLCs (Bairati, 1968; Bertram et al., 1992; Gligorov et al., 2013). These membrane compartments are known to be critical for SC function (Corrigan et al., 2014; Gligorov et al., 2013; Redhai et al., 2016) and are suggested to be involved in the secretory transport route (Corrigan et al., 2014; Gligorov et al., 2013; Redhai et al., 2016). Indeed, Bairati and others have presented evidence indicating that these structures occasionally fuse with the plasma membrane to release their cargo (Bairati, 1968; Corrigan et al., 2014; Redhai et al., 2016). We found that, among the Rab proteins, only Rab 6, 7, 11 and 19 are associated with VLCs. Interestingly, these Rabs define distinct populations of VLCs and form after a maturation process that correlates with the time AGs need to assume their optimal biological functionality (Ruhmann et al., 2016).

Rab6 is a well-studied core Rab protein (Barr, 1999; Deretic and Papermaster, 1993; Iwanami et al., 2016; Satoh et al., 2016; Schotman and Rabouille, 2009) associated with the TGN (Barr, 1999; Martinez et al., 1994) and is reported to regulate retrograde traffic from the Golgi to the ER (Bonifacino and Rojas, 2006; White et al., 1999) and transport destined for secretion (Iwanami et al., 2016; Januschke et al., 2007; Schotman and Rabouille, 2009). In SCs, we found Rab6 associated with the Golgi network as well as a subset of non-Golgi VLCs. The Golgi of the SCs forms an extended central structure, which is unusual for *Drosophila* cells. Most cell types in *Drosophila* possess many solitary Golgi organelles that consist of one *cis*-Golgi and one *trans*-Golgi membrane sheet (Kondylis and Rabouille, 2003, 2009; Rabouille et al., 1999; Ripoche et al., 1994). This organization is viewed as an evolutionary ancestor of the more complex mammalian Golgi-cisternae (Kondylis and Rabouille, 2009). However, the centralization of the Golgi in SCs may indicate a very high membrane traffic turn-over (Liu and Storrie, 2012) and may be consistent with the hypothesis that SCs are involved in the uptake of proteins of foreign origin (Gligorov et al., 2013).

The presence of Rab6 on non-Golgi compartments has also been reported for other secretory cells, like the cells of the SGs (Dunst et al., 2015; Iwanami et al., 2016). Interestingly, in SGs, Rab6 non-Golgi compartments are localized close to the apical membrane and are thought to be involved in the apical secretion of saliva constituents (Dunst et al., 2015; Iwanami et al., 2016). We tested the possibility that Rab6-labelled VLCs are traffic checkpoints for secreted proteins by expressing the published reporter protein Tomato^myr^ in SCs. Consistent with our hypothesis, Tomato^myr^ co-localizes with non-Golgi Rab6-positive VLCs, indicating that Tomato^myr^ is transported via these compartments. However, unlike in the SGs, Rab6 VLCs are not observed in close proximity to the apical plasma membrane, thus, it seems unlikely that the Rab6-positive VLCs are a final secretory compartment before apical secretion. More likely, Rab6 VLCs represent an early compartment on the route towards secretion. Surprisingly, we also found that some Rab6-positive VLCs are also marked by Rab11 domains. Rab11 is another core-Rab protein (Bhuin and Roy, 2014; Hutagalung and Novick, 2011) and has been shown to regulate multiple membrane recycling routes (Casanova et al., 1999; Ullrich et al., 1996). Examining the Tomato^myr^ marker in lines expressing differentially tagged Rab6 and 11, we were able to show the secretion marker in both Rab6 and Rab6/11 VLCs. This is consistent with previous studies, where it was shown that ectopically-expressed Rab11 marks a subset of densely-filled vacuoles in SCs that contain secreted molecules like ANCE (Redhai et al., 2016; Rylett et al., 2007) and DPP (Redhai et al., 2016). Combining these results with our finding that knockdown of *Rab6* in SCs prevents the formation of Rab11 VLCs, we conclude that some Rab11 VLCs are probably downstream compartments involved in the same secretory pathway as Rab6. Lastly, in SCs, we found additional small Rab11-positive (but Rab6-negative) punctae in close proximity to the apical membrane suggesting other Rab11 roles in apical membrane recycling (Casanova et al., 1999; Dunst et al., 2015; Goldenring et al., 1996; Iwanami et al., 2016).

Rab19 is another Rab protein that localizes close to the apical membrane in *Drosophila* SGs (Dunst et al., 2015). Furthermore, *in vitro* interaction experiments have shown that Rab19 can interact with the apical adhesion molecule, Pollux, leading some to suggest a role for rab19 vesicles in apically directed secretion (Dunst et al., 2015; Gillingham et al., 2014; Zhang et al., 1996). Here, we show that Rab19 is strongly expressed and associated with apically localized VLCs, as well as a small portion of the Golgi apparatus. However, we were unable to show that Rab19 plays a role in apical secretion in the SCs. In fact, Rab19 shows no overlap with Rab6-positive membranes and Tomato^myr^ is not trafficked at high levels through Rab19-positive VLCs. Based on these findings, we speculated that Rab19 VLCs are probably not secretory in nature and tested their relation to the Rab7-positive VLCs. Rab7 and Rab19 VLCs appear only in matured SCs and the genetic reduction of Rab7 prevents the formation of Rab19-positive VLCs. By contrast, Rab19 depletion is not sufficient to suppress Rab7 VLCs. This string of evidence suggests that Rab19 VLCs differentiate directly from Rab7 compartments originating from the *cis*-Golgi or endoplasmic reticulum (Bucci et al., 2000; Meresse et al., 1995). An alternative hypothesis that cannot be excluded is that Rab19 VLCs may descend from Rab11 (Rab6-negative) VLCs. To discriminate both possibilities, Rab11 depletion experiments are required. Unfortunately, the currently available tools do not allow such experiments in *Drosophila*. However, the fact that small portions of Rab19 and Rab7 are found within the Golgi organelle and that the appearance of Rab7 VLCs is dependent on Rab6, makes it tempting to speculate that a Rab7>Rab19 endocytic route may be required to recycle proteins originating from paracellular transport (Chan et al., 2011; White et al., 2015).

To assay the biological relevance of the different VLC populations we have assayed the female long-term PMR (Gligorov et al., 2013; Kubli and Bopp, 2012; Ravi Ram and Wolfner, 2007, 2009). The long-term PMR is never developed in females that were mated to mutant males containing SCs without VLCs (Gligorov et al., 2013). As expected for flies with SCs lacking VLCs, the knock down of *Rab6* in SCs prevents the formation of a meaningful long-term PMR of mated females. Interestingly, the knock down of *Rab7*, but not *Rab19* in male SCs also results in the loss of the female long-term PMR. As Rab7 is required for proper Rab19 VLC formation, we originally thought that they would be part of the same pathway, and thus, share the same phenotype. While these two compartments may share some functions, with regards to the long-term PMR, this does not seem to be the case. Based on our results, we believe that the knock down of *Rab7* should block all endocytic traffic, and that this blockage may lead to the loss of long-term PMR through direct or indirect mechanisms. Indeed, this may be the case, as *Rab7* knockdown has been shown to ultimately lead to cell lethality in other systems (Brand and Perrimon, 1993; Chan et al., 2011, 2012; Chinchore et al., 2009). On the other hand, the lack of PMR phenotypes in Rab19 deficient males again confirms that Rab19 is not part of the primary Rab6 VLC secretory traffic route.

We have shown that in male SCs Rab6 is required to establish and maintain two independent trafficking routes. Both transport pathways intersect each other at the Golgi apparatus, but only one, the Rab6/11 VLC branch seems to be essential to deliver factors for the seminal fluid to initiate a lasting PMR in mated females (Corrigan et al., 2014; Redhai et al., 2016). In addition to the work presented here, we have examined the entire Rab machinery in the AG along with a battery of additional protein markers that are accessible through our online resource (https://flyrabag.genev.unige.ch). This work should facilitate future studies on the AG and other studies involving protein transport, paracellular transport and the development/organization of membrane identities.

## Materials and Methods

### Fly stocks

Male collections were performed at 25°C. *D1-Gal4* was generated in the lab (Gligorov et al., 2013), YFP-tagged *rabs* (*Yrabs*) (Dunst et al., 2015), *UAS-Tomato-myristoylation* (Pfeiffer, 2010), *UAS-Lifeactin-Ruby* (Schnorrer, 2011) and *UAS-Golgi-RFP* (Rikhy and Lippincott-Schwartz, 2010) lines were provided by S. Eaton’s laboratory. *UAS-Rab6RNAi* (ID100774) (Keleman et al., 2009; Torres et al., 2014) and *UAS-Rab19RNAi* (ID103653) (Keleman et al., 2009) are available from Vienna *Drosophila* Resource Center and *UAS-Rab7RNAi* line was from M. Gonzalez-Gaitan’s laboratory (University of Geneva) (Assaker et al., 2010; Dickson et al., 2007; Dietzl et al., 2007). Flies were raised at 25°C in tubes on standard yeast-glucose media (8.2% w/w yeast, 8.2% w/w glucose, 1% w/w agar, 1.2% v/w acid mix).

### Immunochemistry

Accessory glands from 5-6 days-old males were dissected in ice-cold Grace’s Insect Medium (BioConcept), fixed for 20 minutes with 4% Formaldehyde (Sigma) at room temperature and stained with one or more of the following antibodies over-night at 4°C: anti-Dlg (Developmental Studies Hybridoma Bank (DHSB)), anti-DE-cadherin (DHSB), or with Phalloidin-546 (Life Technologies). All samples were mounted in Vectashield mounting medium with or without DAPI (Vector Labs). The pictures were taken with a Zeiss LSM700 confocal microscope and evaluated using the FIJI (Schindelin et al., 2012) (Laboratory of Optical and Computational Instrumentation (LOCI), University of Wisconsin-Madison, USA) and IMARIS softwares (Bitplane AG, Zurich, Switzerland).

### Live imaging

Sample were dissected in ice-cold PBS and mounted in PBS onto a coverslip. Samples were imaged at approximatively 20°C by an OMX V3 BLAZE microscope (GE Healthcare Life Sciences; Fig 2). Deconvolution algorithms were applied to the acquired wide-field images using the softWoRx 5.5 software package (GE Healthcare Life Sciences).

### Determination of the distribution of the YRab compartments in the secondary cells

The center of mass of the secondary cells was determined by using Fiji software (a secondary cells was surrounded by using “Freehand selection” and the center of mass was determined by “Measurements”) and a circle of 8.90μm diameter (*ie* average diameter of the apical surface of the secondary cells in contact with the lumen) was drawn by using FIJI software drawing tools; this circle corresponds to the “central” location. The “cortical” and “non-central cytoplasmic” location indicate that compartments are in close proximity to the cellular membrane or not, respectively. The three “Apical” (luminal side), “Medial” and “Basal” (stromal side) portions were determined by counting the number of z-slices covering the secondary cell height and this number was divided by three (Figs 1A-1A’, Fig. S6A).

The expression patterns of the Rab proteins have been described in the SCs from three to seven days-old males. Different terms will be used to describe different Rab-marked structures; we use “vacuole-like compartments” (VLCs) to refer to structures clearly delimited by a fluorescent membrane, whose diameter can vary from 0.3μm to 8μm. The term “small compartments” is used for features >0.5µm, which are homogeneously fluorescent, and “punctate” for distinct structures that are smaller than 0.5μm in diameter. Finally “diffuse” is used for spread out signal without visible particulate structures (Figs 2D, S2B, S6C).

### Receptivity and egg laying assays

New-born virgin males from the different genotypes were put in fresh tubes with dry yeast and stored at 25°C for 5-7 days, 12/12 hours dark/light cycles. The same was done for virgin *Canton-S* (CS) females. On the day before the experiments, fresh tubes containing one virgin female collected 5 days before were set up and kept at 25°C. On the day of mating, one male was added per female-containing tube. For the tubes where mating occurred, the males were removed and the females were kept for receptivity and egg laying assays at 25°C.

Receptivity assay: Mated females were put in fresh tubes and 4 days after mating, one CS male was added into the tube. The tubes where mating occurs were counted, while for the tubes where the flies did not copulate, the males were removed and the tubes were kept for the next receptivity assay *ie* 10 days after the initial mating (6 days later).

Egg laying assay: single females were transferred every day in a fresh tube and the eggs laid were counted (over a period of 10 days).

## Acknowledgments

We thank Dr. Marko Brankatschk and Prof. Suzanne Eaton for welcoming us into their laboratory and sharing the fly Rab library. We thank Dr. Siamak Redhai and Dr. Ian Dobbie (MICRON, Oxford) for help with live imaging of accessory glands. We thank Dr. Virginie Sabado and the members of Pr. Emi Nagoshi’s laboratory for technical advice on sample preparation, microscopy and fly behavioral analysis. We thank the Bioimaging Center of the University of Geneva and the Micron Oxford Advanced Bioimaging Unit for microscopy facilities, Dr. Marko Brankatschk, the Karch’s laboratory, Prof. Mariana Wolfner, Prof. Marcos Gonzalez-Gaitan and Prof. Jean Gruenberg for advice and support regarding this project. And finally we thank Dr. Dragan Gligorov for initiating this research project.

This work was supported by grants from the Donation Claraz, the State of Geneva, the Swiss National Fund for Research to FK, the BBSRC (BB/K017462/1) and the the Wellcome Trust (Strategic Award #091911 and 102347/Z/13/Z)

